# Identification of novel genetic regions associated with resistance to European canker in apple

**DOI:** 10.1101/2021.12.20.473552

**Authors:** Amanda Karlström, Antonio Gómez-Cortecero, Charlotte F Nellist, Matthew Ordidge, Jim M. Dunwell, Richard J Harrison

## Abstract

Resistance to *Neonectria ditissima*, the fungus causing European canker in apple, was studied in a multiparental population of apple scions using several phenotyping methods. The studied population consists of individuals from multiple families connected through a common pedigree. The degree of disease of each individual in the population was assessed in three experiments: artificial inoculations of detached dormant shoots, potted trees in a glasshouse and in a replicated field experiment. The genetic basis of the differences in disease was studied using a pedigree-based analysis (PBA). Three quantitative trait loci (QTL), on linkage groups (LG) 6, 8 and 10 were identified in more than one of the phenotyping strategies. An additional four QTL, on LG 2, 5, 15 and 16 were only identified in the field experiment. The QTL on LG2 and 16 were further validated in a biparental population. QTL effect sizes were small to moderate with 4.3 to 19 % of variance explained by a single QTL. A subsequent analysis of QTL haplotypes revealed a dynamic response to this disease, in which the estimated effect of a haplotype varied over the field time-points. Two groups of QTL-haplotypes could be distinguished, one that displayed increased effect and one with a constant effect across time-points. These results suggest that there are different modes of control of *N. ditissima* in the early stages of infection compared to later time-points of disease development. It also shows that multiple QTL will need to be considered to improve resistance to European canker in apple breeding germplasm.

## Introduction

European apple canker, caused by the fungal pathogen *Neonectria ditissima*, infects a wide range of hosts, including apple (*Malus spp*.), pear (*Pyrus spp*.) and a range of other broad-leaved perennial species (Walter, et al., 2015). The disease is widespread in apple orchards in regions with temperate and wet climates (Beresford & Kim, 2011; Weber, 2014).

*N. ditissima* is foremost a wood pathogen, which enters the host through natural or artificial wounds such as leaf scars, pruning cuts or lenticels and cracks in the bark (Weber, 2014). The disease symptoms are trunk cankers, branch lesions and die-back. In severe cases the lesions expand to girdle the whole main trunk of the tree and kill all branches above the point of the canker (Ghasemkhani, 2015).

European canke control strategies are removal of infected wood through pruning and the limitation of new infections through the application of fungicides when wound incidence is high (Walter et al., 2015; Walter et al., 2017). Genetic variation in *N. ditissima* resistance has been documented in commercial apple varieties of *Malus x domestica*, wild species of *Malus* as well as in apple rootstocks (Garkava-Gustavsson et al., 2013; Ghasemkhani, 2015; Gómez-Cortecero et al., 2016; Van de Weg, 1989a, 1992). Previous studies of the inheritance of canker resistance in apple progenies demonstrate that the resistance is inherited quantitatively (Gómez-Cortecero et al., 2016), though QTLs of major effect might be present too (Van de Weg, 1989b). The relative levels of resistance of mature trees of parental cultivars correspond well with the relative levels of juvenile full-sib families, indicating that juvenile material may be used in inheritance studies (Van de Weg 1989b). The crab apple accession *Malus robusta* ‘Robusta 5’ has shown tolerance to infection by *N. ditissima* in multiple studies (Bus et al., 2010; Gómez-Cortecero et al., 2016). Bus et al. (2019) genetically mapped a QTL for disease incidence from ‘Robusta 5’ in a bi-parental cross between the apple rootstock ‘M9’ x ‘Robusta 5’. In the study, a single medium effect size QTL on linkage group 14 was identified by mapping the disease incidence in segregating progeny genotyped with simple-sequence repeats (SSR)-markers. This QTL accounted for around 40% of the variance in disease, but the effect size was dependent upon the phenotyping method (Bus et al., 2019). Although there are commercial scion cultivars tolerant to apple canker, there is no published information on which chromosomal regions control the quantitative resistance found in this germplasm.

Response of plants to disease has shown temporal variation in several plant-microbe interactions (Chang et al., 2018; Li et al., 2012; Li et al., 2007; Mohler & Stadlmeier, 2019; Welz et al., 1999). The dynamic QTL methods in these studies analyse the phenotypic variation at different times during infection, whereas conventional methods would analyse the cumulative disease phenotype at the end. In these studies, resistance QTL were uniquely identified at different stages of disease or plant development, indicating that the genes controlling the response to disease have temporal expression patterns. This type of, so called, dynamic response has not been reported for interactions with wood pathogens.

In the present study, the genetic basis of resistance to *N. ditissima* in apple scion germplasm was studied through QTL mapping in a multiparental population using a pedigree-based Bayesian analysis (Bink et al., 2014). Three phenotyping strategies (field, potted trees and shoots) were used to determine whether rapid phenotyping methods can replace field experiments for QTL discovery.

## Results

### Phenotypic analysis

The multiparental population showed a normal distribution of disease levels to European canker in all experiments (Suppl. Fig. 1A-H). Mean values for each family and European canker phenotype are shown in Supplementary Table 1. A comparison of phenotypes for parental apple genotypes and a standard set of cultivars with known resistance to European canker is shown in Supplementary Figure 1.

All genotypes exhibited disease symptoms upon infection, hence there was no evidence of complete resistance to this disease in the genotypes studied. The overall infection success of the artificial inoculations was >95 % in all three phenotyping experiments.

The broad sense heritability (H^2^) from the different phenotyping events and measurements is shown in Supplementary table 2 and correlations between these are shown in Figure 1. The canker phenotypes recorded from detached shoots and potted trees resulted in lower estimates of heritability than data from the field experiment, 0.16, 0.46 and 0.54-0.76, respectively (Suppl. Table 2).

**Figure 1.**
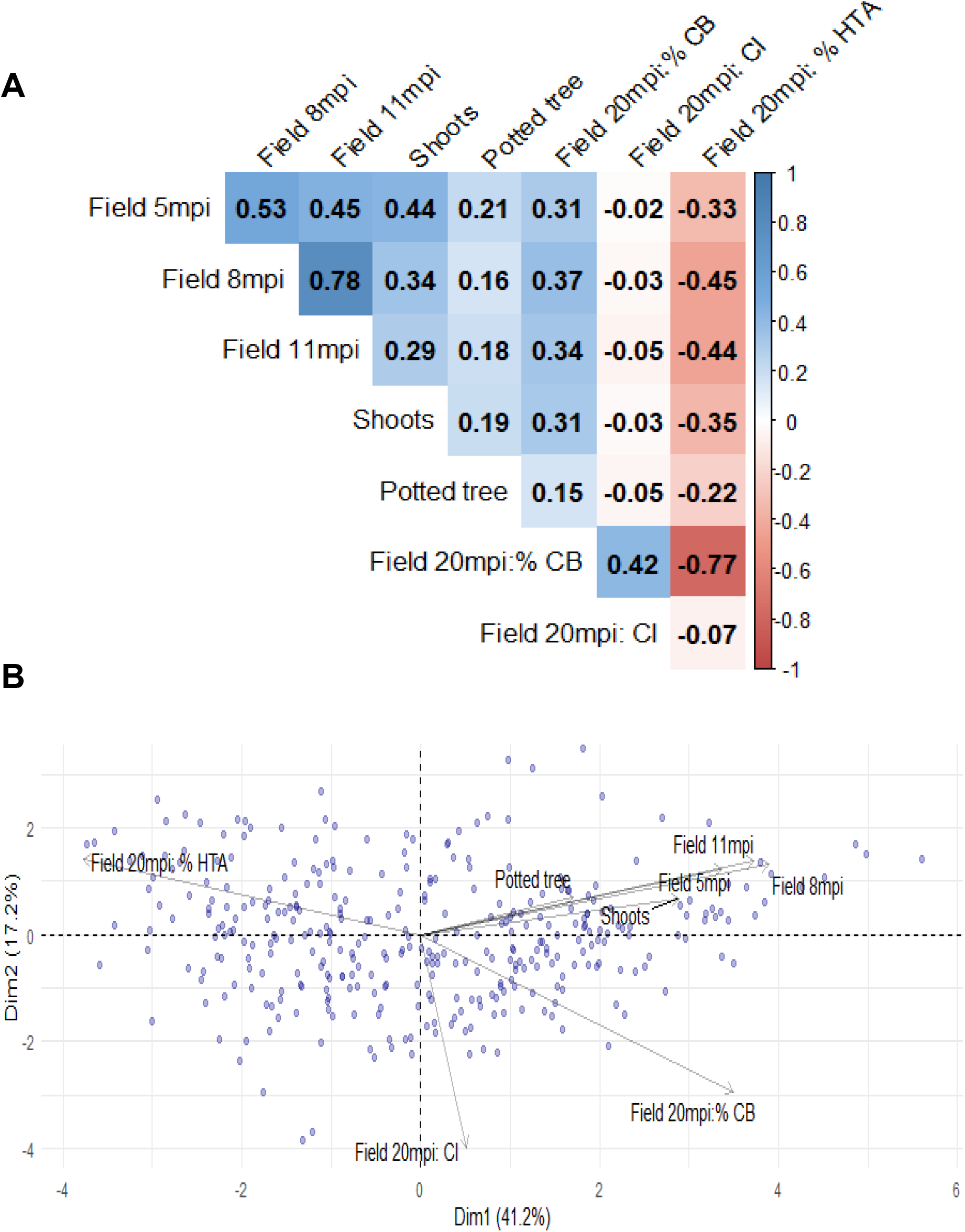
Correlation matrix and principal component analysis (PCA) of phenotypic susceptibility data to European canker in a multiparental population of apple. (a) Correlation matrix showing Pearson’s correlation coefficient of phenotype data. (b) PCA biplot for the first and second component. The length of the arrows approximates the variance of the variables, whereas the angles between them (cosine) approximate their correlations.

Within the field experiment, the Pearson correlation for genotypic Best Linear Unbiased Estimates (BLUEs) of canker lesion size was relatively low between 5 and 11 months post inoculation (mpi, r= 0.45, *p* < 0.001). The highest correlation was observed between the two later time-points (8 and 11 mpi, (*r=* 0.78, *p < 0*.*001*). The BLUEs from the potted tree experiment had a low correlation with the susceptibility in the other phenotyping experiments (see Fig. 1). The shoot experiment had a higher correlation to the early stage of field infection at 5 mpi (*r=* 0.44, *p < 0*.*001*), compared to the two later time-points (Fig. 1A). Canker Index (CI) had a moderate correlation to percent cankered branches (%CB, *r=* 0.42, *p < 0*.*001*) but was not correlated to the other phenotypes. The relationship between different European canker phenotypes is also visualised in the biplot for the correlation matrix PCA (Fig. 1B). The first and second principal component from the PCA explained 41.6 and 17.3 % of the variance, respectively (Suppl. Fig. 3). The loadings from field 5, 8, 11 mpi, shoot and potted tree experiment are clustered together, indicating a high degree of correlation between these variables for the two first principal components. Loadings from these variables also show a positive correlation to PC1 (*r=*0.19-0.43), whereas percent healthy tree area (%HTA) is negatively correlated to the same PC (*r=*-0.42). Nevertheless, the potted tree data had the highest correlation (*r=*0.84) to PC3, to which 12% of the total variation was attributed. The CI loading was largely uncorrelated to PC1 (*r=*0.05) but had a strong negative correlation to PC2 (*r=*-0.70).

A correlation between %HTA and the total number of branches per tree was observed (*r*=0.27, data not shown), suggesting that more vigorous trees were able to survive infection and maintain foliage. A similar negative correlation was found for %CB (*r*=-28). Surprisingly, there was also a weak positive correlation between 5 mpi and number of branches (*r=*0.17).

### Identification of QTL associated with resistance

The Bayesian QTL mapping analysis revealed a total of seven linkage groups involved in the response to European canker (Fig. 2 and Table 1). There was positive (2lnBF>2) or strong evidence (2lnBF>5) for a QTL in more than one phenotyping experiment for three of the LGs: 6, 8 and 10. For the remaining four LGs there was strong evidence for a QTL in one of the phenotyping events at the linkage group level. The variance attributed to individual QTL was small to moderate with effect sizes of 5 to 19 %.

**Figure 2.**
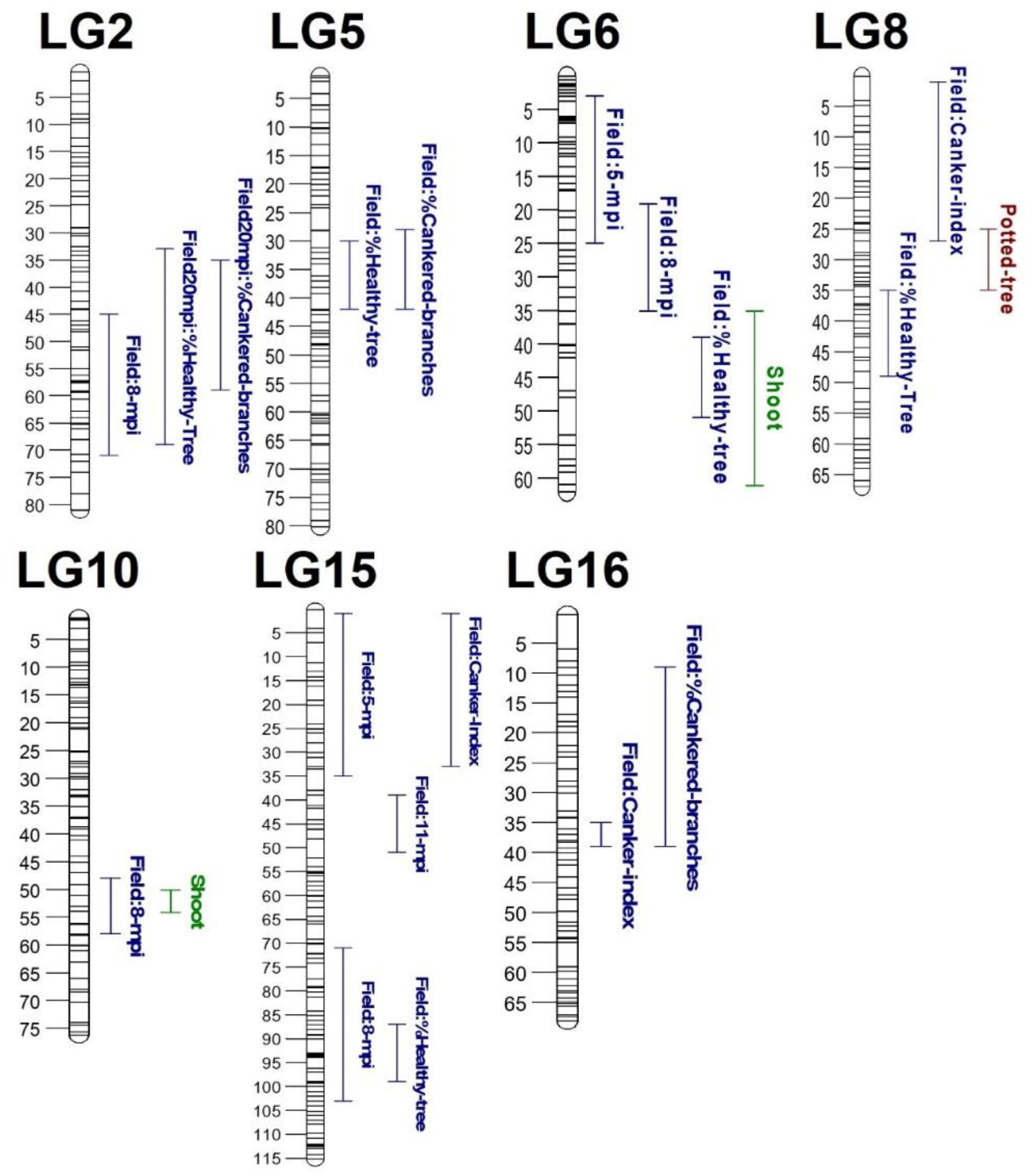
QTL regions identified to be associated with resistance to European apple canker. The figure shows QTL intervals from each measured phenotype. Different colours indicate distinct phenotyping experiments.

**Table 1.**
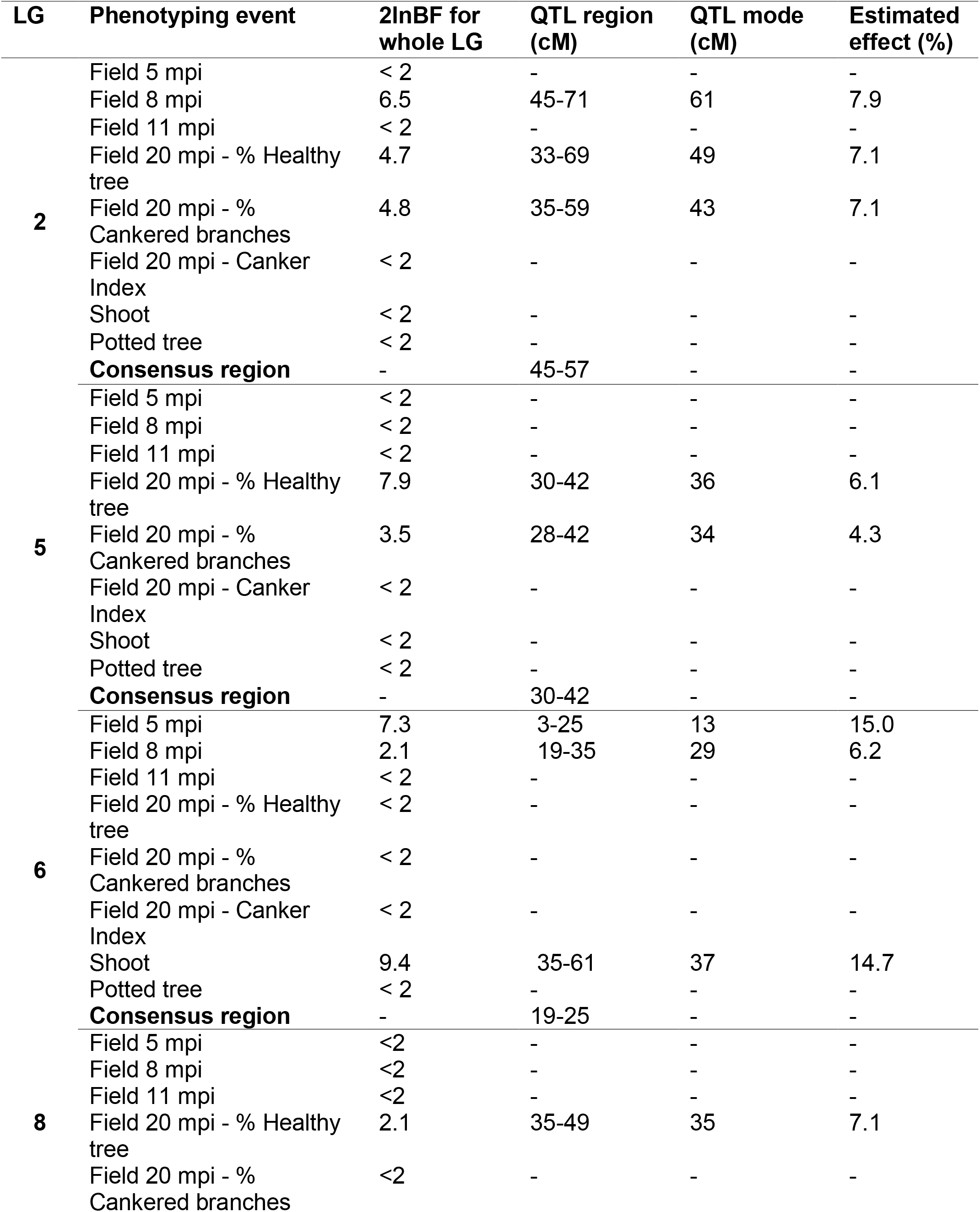
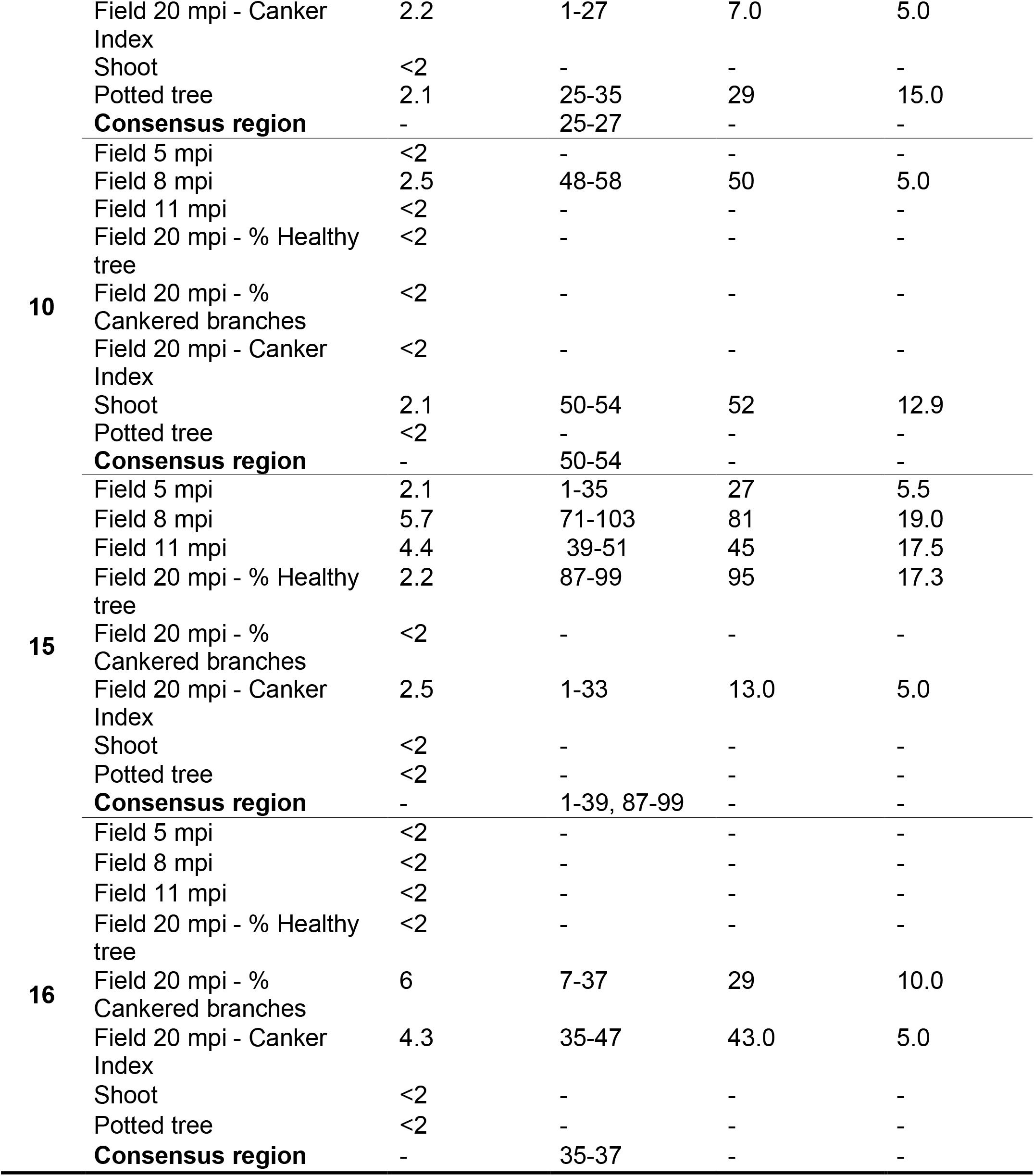
Summary of the results from the quantitative trait loci (QTL) mapping of resistance to European canker with the FlexQTL software. QTL regions reported consist of successive 2-cM bins with 2ln Bayes factors greater than 2.

There was strong evidence for a QTL on LG6 in both the shoot experiment and at 5 mpi in the field (2lnBF of 9.4 and 7.3, respectively). There was also positive evidence (2lnBF⋝2) for a QTL on this linkage group at 8 mpi (Table 1). However, two separate QTL regions on LG6 were indicated, depending on the analysed phenotype (Fig. 2). There was no positive evidence for the presence of more than one QTL in the output from FlexQTL. The FlexQTL analysis showed that QTL on LG6 had a larger effect at the earlier stage of field infection, as the QTL explained 15 % of the variation in shoots and at 5 mpi, but less than 7 % at later field time-points.

A QTL on LG8 was identified in data for %HTA (2lnBF=2.1), CI (2lnBF=5.6), and the potted tree experiment (2lnBF=2.1). Hence, this QTL was found in more than one type of experiment. Depending on phenotype, the variance attributed to this QTL ranged between 9.3-15%. A third QTL, located on LG10, was positively identified using two types of phenotyping methods. There was positive evidence for this QTL in the field 8 mpi and in the shoot experiment (2lnBF=2.5 and 2.1, respectively).

A QTL on LG5 was positively identified in data from the field experiment at 20 mpi but was not identified at earlier time-points nor in the other phenotyping experiments. There was strong evidence (2lnBF = 7.9) for a QTL on LG5 for %HTA and positive evidence (2lnBF = 4.4) for %CB. Hence, this QTL was only identified after a prolonged period of infection.

The variance attributed to QTL on LG15 was the highest among all discovered QTL, with an effect size of 19 and 17.5 % at 8 and 11 mpi respectively. Nevertheless, the effect of this QTL was much lower at 5 mpi (5.5%). Three regions on LG15 were identified to have an association with the susceptibility to canker depending on phenotype (Fig. 2). However, there was no positive evidence for the presence of more than one QTL in the analysis.

A further two QTL were identified on LG2 and LG16. The QTL analysis did not discern any obvious temporal patterns in effect sizes for these QTL.

### Haplotype analysis

A haplotype analysis was conducted to understand which haplotypes contributed to resistance to *N. ditissima* at different stages of infection. The effect of all haploblocks within the QTL regions was tested on lesion size data at 5, 8 and 11 mpi in the field. The haploblocks (HB) with the most significant effect on canker lesion size from each QTL region and the number of unique haplotypes within the multiparental population are shown in Supplementary Table 3.

Figure 3 shows the estimated percent deviation from the mean of resistant and susceptible/neutral haploblock alleles for the three time-points. Haplotypes were only included if they were present in a parent segregating for European canker resistance at that QTL locus, as haplotype effects from non-segregating parents could not be reliably estimated. A total of 33 haploblock alleles were present in segregating parents across all seven genetic regions.

**Figure 3.**
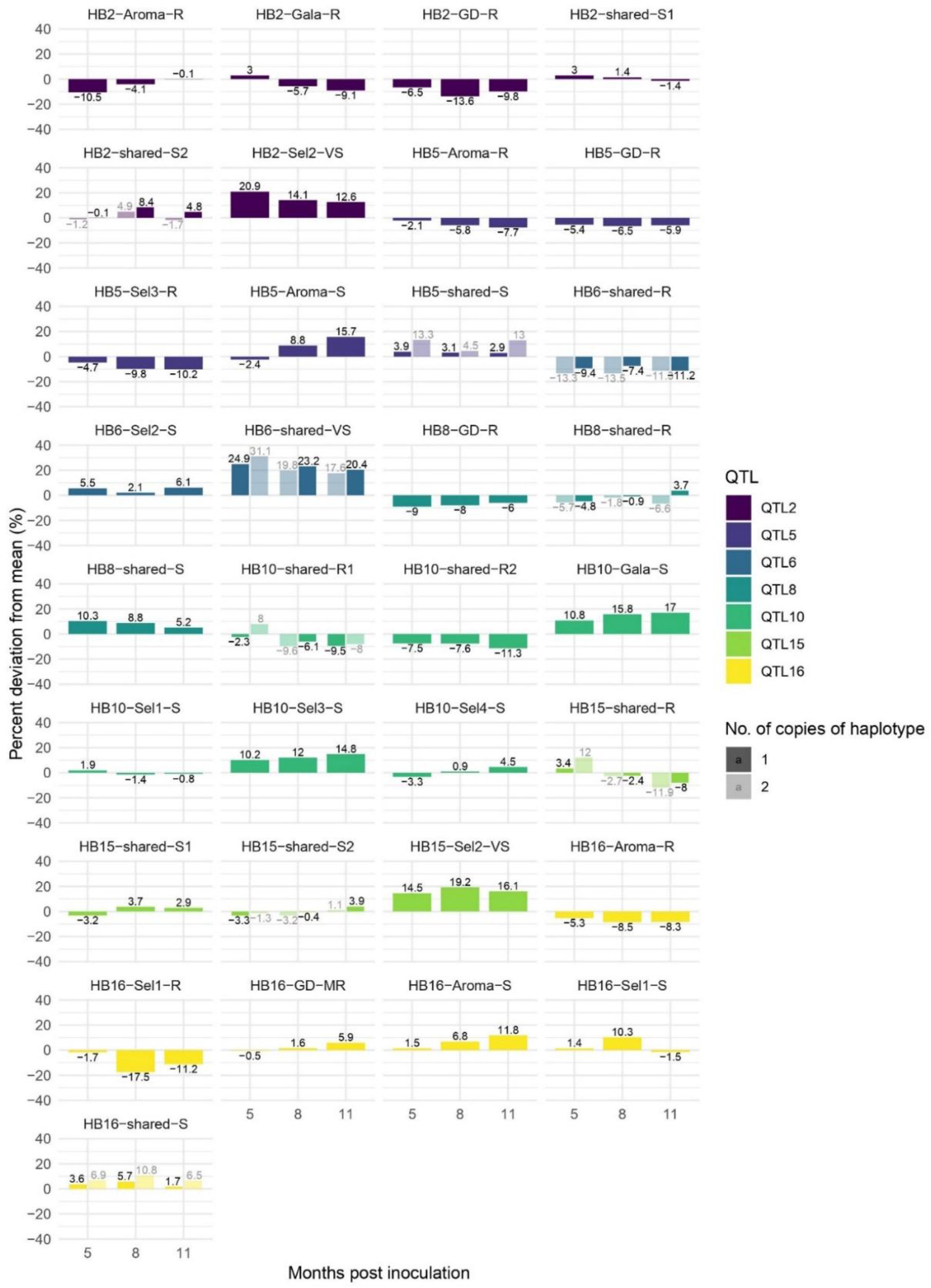
Estimated percent deviation from mean lesion size at 5, 8 and 11 months post inoculation for individuals with 1 or 2 copies of haplotypes. The haplotypes shown are present in at least one parent that segregates for resistance to European apple canker at the QTL-locus. Haplotypes denoted R have a resistant effect while S and VS have susceptible and very susceptible effects.

The estimated effects of some haplotypes varied over the three time-points (Fig. 3). For HB5, the percent deviation from the mean more than doubled over the assessed time-period for two out of three resistant haplotypes (Fig. 3). Furthermore, the allele HB15-shared-R2 had a susceptible effect on lesion size at 5 mpi but a resistant effect at 11 mpi, with -8 and -11 % deviation from mean for individuals with one or two copies, respectively, of the haplotype. The resistant alleles HB16-Aroma-R and HB16-Sel1-R also showed increased effects on resistance with time.

The haplotype alleles with the largest negative effects on lesion size were found within HB2, HB6, HB15 and HB16 (Fig. 3). The effect of HB6-shared-R, which ranged between -7.4 to -13.5 %, was similar across time-points and for individuals carrying one or two copies of the allele. This haplotype was inherited from four of the parents, Gala, Golden Delicious, EM-Selection-2 and EM-Selection-4 and could be traced back to the founding cultivars ‘Grimes Golden’ and ‘Jonathan’. These two founders were also the origin of the haplotype HB15-shared-R. The resistant effect of HB15-shared-R increased with time and was estimated to reduce lesion size with -8 and -12 % at 11 mpi for individuals with one or two copies, respectively, of the haplotype. HB15-shared-R segregated in two of the parents, Golden Delicious and EM-Selection-4. This haplotype was also present in a third parent (EM-Selection-1), which had two alleles with a resistant effect on lesion size at this locus.

Three haploblock alleles were associated with large increases in lesion size: HB2-Sel2-VS, HB6-shared-VS and HB15-Sel3-VS (Fig. 3). Individuals heterozygous for HB2-Sel2-VS had an estimated +13-21 % increase in lesion size, depending on time-point. The origin of this haplotype could not be traced further back than to the unreleased selection ‘EM002’ due to the lack of available material in the germplasm collection (Fig. 5). HB6-shared-VS was associated with an up to +31% increase in lesion size compared to the mean (Fig. 3). The allele was inherited from the three parents Gala, EM-Selection-3 and EM-Selection-4, of which EM-Selection-3 was homozygous for the haplotype. Hence, HB6-shared-VS only segregated in the family MDX061. This allele was present in many of the founding cultivars, including ‘Delicious’, ‘Jonathan’, and ‘Ingrid-Marie’.

### Validation of haplotype effects in a biparental population

A significant haplotype-trait association was identified for two of the seven QTL regions in progeny from a ‘Golden Delicious’ x ‘M9’cross. Based on the results from the multiparental population, QTL on five LG were expected to segregate in ‘Golden Delicious’: LG 2, 5, 8, 15 and 16. The segregation of resistance loci in ‘M9’ was not previously known. Mean AUDPC and disease distribution for haploblock alleles for which ‘Golden Delicious’ segregates are shown in supplementary table 5 and Fig. 4, respectively.

**Figure 4.**
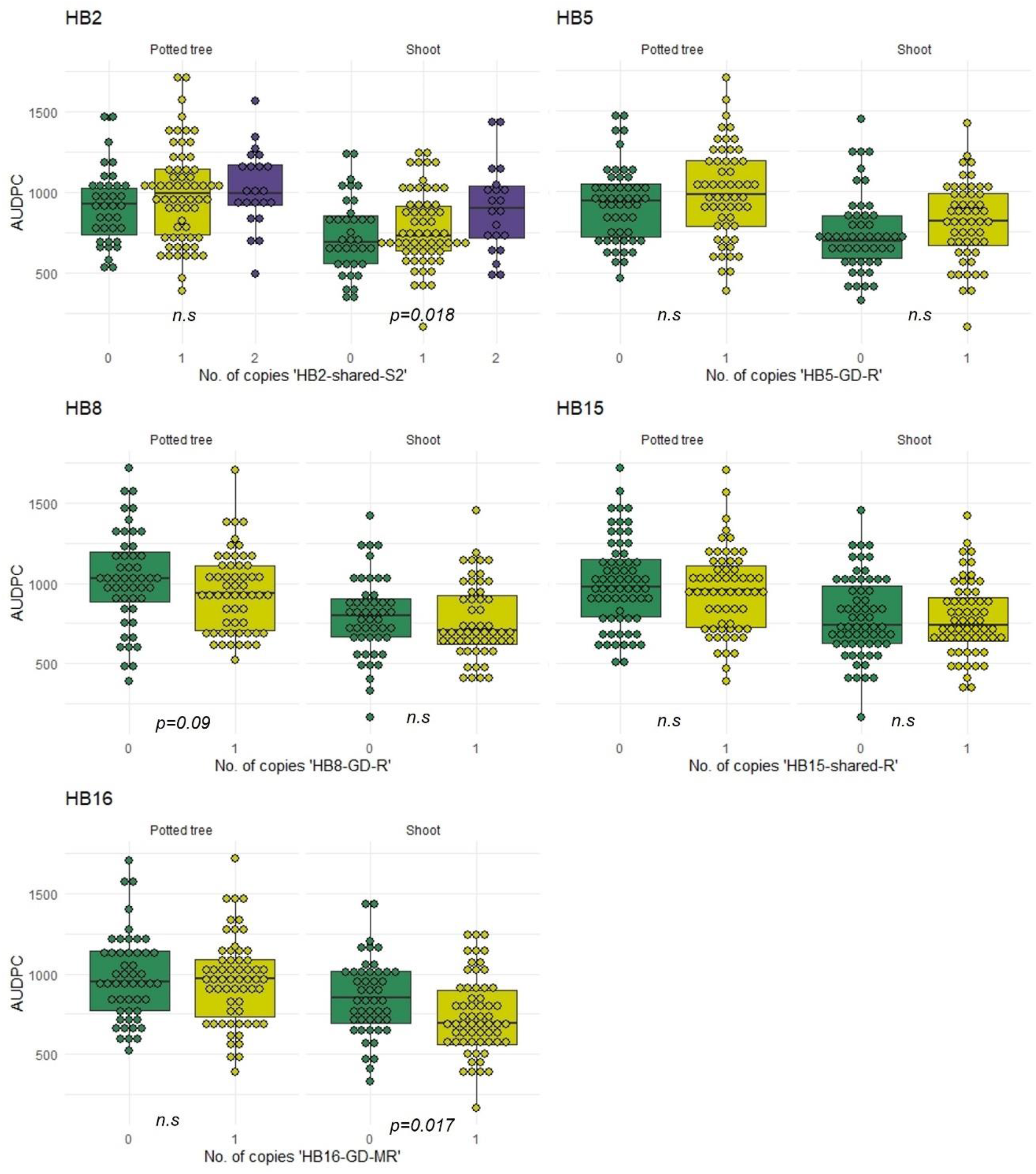
Validation of haploblocks for European canker resistance in two phenotyping experiments. Boxplots show the Area Under Disease Progress Curve (AUDPC) for individuals from a biparental cross between ‘Golden Delicious’ x ‘M9’.

The haplotype ‘HB2-shared-S2’ was inherited from both parents and was associated with an increase in AUDPC (Fig. 4). There was a significant effect of this haplotype (*p=*0.02) in data from the detached shoot experiment. Additionally, the haplotype ‘HB16-GD-MR’ was confirmed to be associated with resistance to European canker in the detached shoot experiment (*p=*0.017). Individuals with one copy of this allele had a reduction in mean AUDPC with 28 and 118 units in the potted tree and shoot experiment, respectively. No haploblock alleles had a significant effect on AUDPC in the potted tree experiment.

Progeny with the allele ‘HB8-GD-R’ had a reduced mean AUDPC with 94, and 22 units, respectively, in the potted tree and shoot experiment. Nevertheless, the effect of this haplotype was non-significant (*p=*0.058). Surprisingly, progeny with the haplotype ‘HB5-GD-R’ had a higher mean AUDPC compared to individuals that inherited the susceptible allele ‘HB6-shared-S’ in both experiments (Suppl. Table 5).

The proportion of variation attributed to significant haplotypes was low, with 4.5-5.4% variance explained by single haplotypes in the detached shoot experiment. The total variance due to significant haplotypes was 10% in this experiment.

## Discussion

### Identified resistance loci widely distributed in apple scion germplasm

Seven linkage groups were shown to be associated with the level of resistance to European apple canker in a multiparental scion population. Three of these QTL were identified in more than one experiment using the same population: the QTLs on LG6, LG8 and LG10. Furthermore, the effect of the most significant haploblocks from two of the QTL-regions (HB2 and HB16) were validated in separate experiments with a biparental population derived from a cross between ‘Golden Delicious’ and ‘M9’. Thus, five out of the seven QTL-regions were confirmed in more than one experiment. All the identified QTL had small to moderate effects on disease expression. The effects of the identified QTL exhibited two different patterns: increased effect with time and stable effects across time-points. The phenotypes from the potted tree and shoot experiment were considered to mimic early-stage disease responses, as these experiments ended within a few months after commencement.

This study reveals that resistance loci to European apple canker are present in apple germplasm commonly found as parents and grandparents in modern breeding programmes. Four haplotypes, HB2-GD-R, HB6-shared-R, HB10-shared-R1 and HB15-shared-R, with the largest resistance effects, are all present in ‘Golden Delicious’, a variety that has been reported to feature in the pedigree of 51% out of 500 modern apple varieties with Central-European and US origin (Bannier, 2011). However, the resistance loci on LG6 and LG10 from ‘Golden Delicious’ could not be identified in a bi-parental population derived from this variety as it does not segregate at these positions. Resistant haplotypes described in this study are also shared by varieties that have been reported to be moderately tolerant to *N. ditissima* in previous studies, such as ‘Santana’ (HB 2, 6, 10), ‘Priscilla’ (HB 2 and 6), ‘Elstar’ (HB 2, 6, 10) and ‘Jonathan’ (HB 6, 10, 15 and 16) (Amponsah et al., 2017; Van de Weg, 1989; Wenneker et al., 2017). Several of the identified resistant haplotypes in this study were inherited from the Swedish variety ‘Aroma’, which has been reported to show a medium level of resistance (Garkava-Gustavsson et al., 2013; Gomez-Cortecero et al., 2016). These haplotypes are derived from the founding parent ‘Filippa’ and unlikely to be widespread in germplasm outside of Scandinavia.

Interestingly, the QTL-region on LG15 has also been shown to be associated with increased resistance to the fungal wood pathogen *Valsa mali* (Tan et al., 2017). *In the study, the resistant locus was mapped on LG15 of the parent ‘Jonathan’, a variety that is also predicted by us to segregate for resistance to N. ditissima* at this locus. This suggests that this locus may have a wider role in the resistance to wood pathogens in apple.

### Evidence of dynamic QTL effects in resistance to European apple canker

Disease development after infection by *N. ditissima* is a slow process, often characterised by a prolonged symptomless period followed by necrotic lesions that spread within the bark. During the initial stage of infection there is no macroscopic visible resistance response from the host tree; however, as time progresses, some host genotypes develop callus around the boundaries of the canker lesion (Weber, 2014). By mapping lesion size data from each time-point, we were able to study the dynamics of genetic effects during *N. ditissima* colonisation. This showed a temporal pattern in effects of QTL and its related SNP haplotypes. The distinction between early and late response is supported by the correlation of disease phenotypes, with a higher correlation between the detached shoot, potted tree and 5 mpi in the field compared to phenotypes from the later field time-points.

The results show evidence of two QTL, on LG5 and LG16, with larger genetic effects at the later stages of disease development. These later responses may be involved in cell-wall modifications and lignin deposition (Lee et al., 2019; Lionetti et al., 2017; Xie et al., 2018; Yan et al., 2019) and might be expected to be conferred by different groups of genes. The QTL-region on LG5 contained genes with a predicted function in cell wall modification; namely two *CELLULOSE SYNTHASE A* genes. These have been shown to be specific to cellulose biosynthesis in secondary xylem cells in poplar, and could play a role in the metabolic coordination of cellulose and lignin biosynthesis during *N. ditissima* infection (Wu et al., 2000).

### Phenotyping methods

This study involved replicated assessments of resistance to European canker using three types of plant material (dormant shoots, actively growing trees in pots and field planted trees) and four types of phenotypes (lesion size, healthy tree area, percent branches with canker and number of canker lesions). It was evident from the PCA biplot (Figure 1b) that there was a positive correlation between lesion size, irrespective of plant material used. As the two first components in the PCA explained most of the variation (59%), it can be assumed that these methods assess similar types of pathogen responses. This is supported by the evidence for a QTL on LG6 and LG10 in both canker lesion data from the field and the shoot experiment. Nevertheless, only three out of seven QTL could be positively identified in more than one experiment. This may be due to the relatively small effect sizes of identified QTL, which has the implication that they sometimes fall below the threshold of detection (Würschum, 2012). The low heritability of the detached shoot experiment (H^2^=0.16), compared to the other phenotyping methods, indicates a large degree of variation between replicates within this experiment. This is in concordance with previous studies using this type of phenotyping assay (Garkava-Gustafsson et al., 2016; Scheper et al., 2018). Data from the potted tree experiment only revealed one QTL with positive evidence and had a relatively low correlation to the other canker lesion phenotypes, despite reports that similar experiments could provide sufficient resolution to differentiate between susceptible and resistant varieties (Garkava-Gustafsson et al., 2016; Wenneker et al., 2017). One influencing factor could be the timing of inoculations, as the potted trees in our experiment were inoculated while actively growing, whereas the other studies inoculated closer to leaf-fall. Generally, it can be concluded that long term experiments are needed to fully capture all resistance responses to European canker in apple and rapid phenotyping methods are therefore not able to replace field experiments.

The canker index (CI) used in this study provides information on the number of secondary canker lesions that had developed 20 mpi. The phenotype correlations and PCA indicate that CI is largely uncorrelated to all other phenotypic traits except for %CB. This would suggest that infection and colonization provide information on different components of resistance. This finding is in accordance with research by Garkava-Gustafsson et al. (2016) but contradicts results from by Wenneker et al. (2017), where moderate-strong correlations were found between infection percentage and colonization. The discrepancies between the different studies might be due to differences in experimental approaches, and neither of the other studies used trees which had already been infected with canker.

The correlation observed between %HTA and %CB and total number of tree branches suggest a relationship between tree vigour and canker tolerance. The identification of a QTL on LG5, which was only revealed from %HTA and %CB data, could therefore be influenced by tree growth. Indeed, this linkage group has previously been associated to tree growth traits in apple (Harrison et al., 2016; Kenis and Keulemans, 2007). Nonetheless, there was a significant effect of individual haplotypes from this QTL region on canker lesion size at 11 mpi, indicating a direct effect on pathogen colonization.

### Multi-QTL approach required to breed for improved resistance

The results from this study show that QTL present in commonly used apple breeding germplasm have a low to medium effect on resistance to *N. ditissima*. Hence, multiple QTL will need to be considered to improve resistance through breeding. As phenotyping of resistance to European apple canker is time-consuming and costly, marker-assisted selection would greatly benefit the selection process. Genomic prediction uses methods of simultaneously estimating the effect of a large set of markers distributed across the genome, and would therefore provide a good alternative for the selection of a multi-QTL trait such as resistance to European apple canker (Crossa et al., 2017) Medium effect apple canker QTL could be incorporated in the genomic prediction model to achieve higher prediction accuracies for this trait (Lee et al., 2019).

## Materials and Methods

### Plant material

A multiparental population, comprising 317 individuals from four full-sib and one half-sib family and their parents were used for this study (Fig. 5, Supp. Table 1). The families were chosen based on a subset of progeny from each family showing segregation for resistance to European apple canker (Gómez-Cortecero et al., 2016). The experiments were carried out while the trees were juvenile.

**Figure 5.**
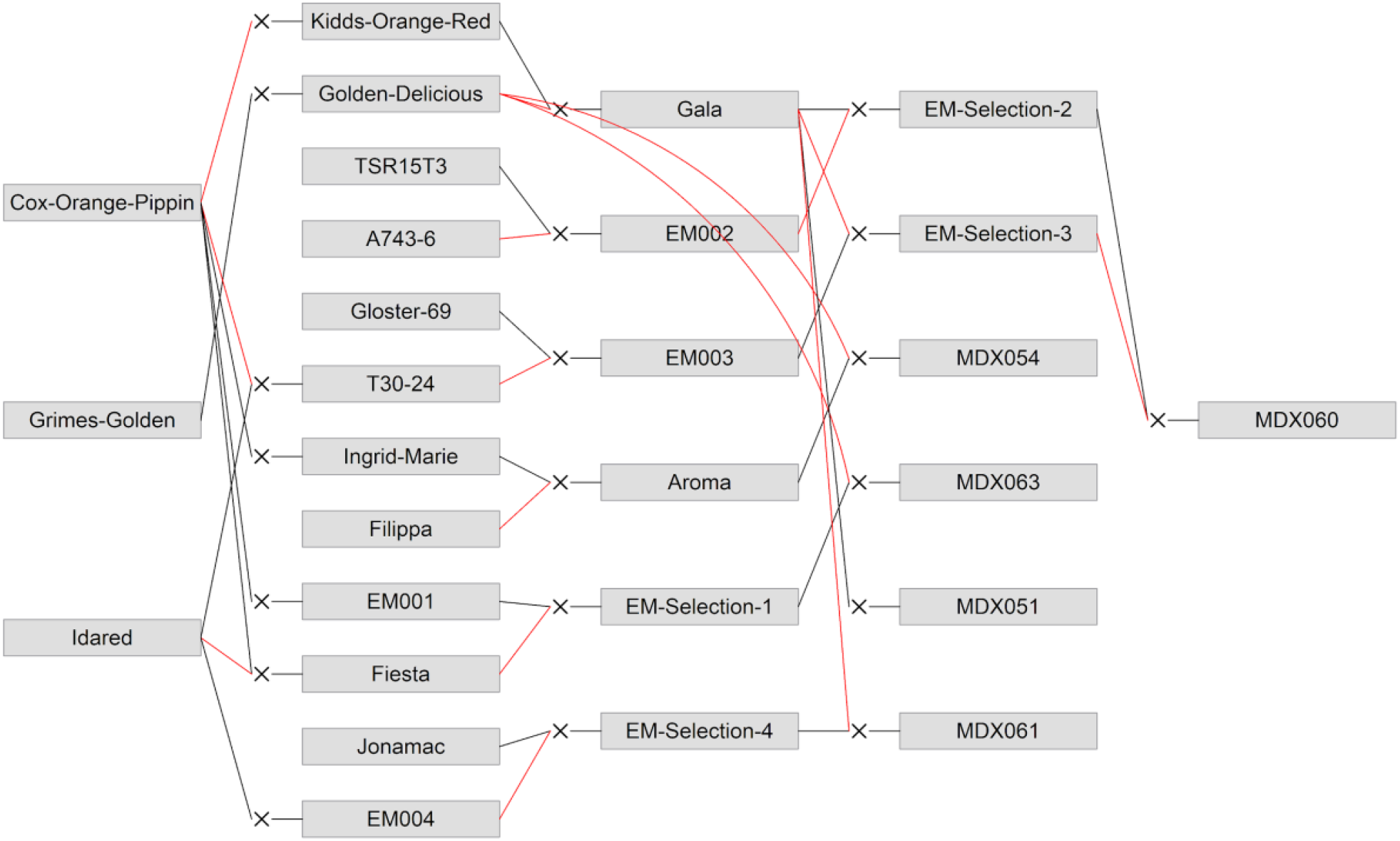
Pedigree showing the parents and progenitors of the five families included in this study; MDX051, MDX054, MDX060, MDX061 and MDX063. Black lines indicate the maternal and red lines the paternal parent.

The phenotyping of resistance to *N. ditissima* was performed by carrying out three types of experiments, namely, artificial inoculations of field planted trees, potted trees in the glasshouse and detached shoots.

Seven replicate grafts were made from each member of the population, along with the parental genotypes. All genotypes were grafted onto ‘M9’ EMLA rootstocks. Grafted plants were grown on in 2L pots in polytunnels before planting out four plants for field experiments (after nine months) and including three plants in potted tree experiments (after seven months).

### *Neonectria ditissima* inoculum

*N. ditissima* isolate Hg199 was used for all pathogenicity screens. Hg199 has been shown to be highly pathogenic on apple (Gómez-Cortecero et al., 2016). Inoculum for each pathogenicity screen was prepared according to Gómez-Cortecero et al. (2016). A concentration of 10^5^ macroconidia/ml was used in all experiments. Inoculations that did not result in lesions were handled as missing data.

## Resistance phenotyping

### Field experiment

The field experiment was conducted in a randomised block design in a field in East Malling, UK. The varieties ‘Cox Orange Pippin’ (susceptible), ‘Jonathan’ (medium resistant), and ‘Santana’ (medium resistant) were included in the experiments as references.

Artificial leaf scar inoculations were conducted following twelve months of establishment in November, 2018. Inoculations were conducted as per Gómez-Cortecero et al. (2016) with a few modifications. Five artificial leaf scars were inoculated per tree, with each leaf scar positioned on a separate branch. Individual inoculations within a tree were considered as pseudo-replicates; each was marked to allow repeated measurements. An inoculum volume of 6 μl at was pipetted onto each artificial leaf-scar wound and covered with petroleum jelly (Vaseline) after absorption. Each pseudo-replicate within a block was inoculated with the same source of inoculum. Conidial germination rate was determined as per Walter et al. (2016) and ranged between 50-86% for different inoculation days.

Canker lesion development was measured with digital calipers at three time-points, five, eight and eleven mpi. Lesions that reached the full length of the branch were recorded as missing data. A final assessment of the field experiment was conducted at 20 mpi, in which the following three phenotypes were recorded; percent branch area with foliage (healthy tree area; %HTA), percentage of all branches with cankers (cankered branches, %CB), and number of cankers. The number of cankers included the sites of inoculation as well as secondary canker lesions on the tree. A canker index (CI) was thereafter calculated by dividing number of cankers with the number of branches on each tree. Trees that were completely covered with canker were removed from the canker index.

### Potted tree experiment

The experiment was conducted between August-November of 2018. Potted trees were placed in glasshouse compartments with misting lines set to maintain a relative humidity above 80%. Trees were drip-irrigated throughout the experiment. Glasshouse compartments were equipped with supplementary lighting (SON-T 400w) set at 16:8 light:dark hours. A chilling-fan ensured a maximum temperature of 22°C.

Two artificial leaf scar wounds (pseudo-replicates) were inoculated on the main leader of each tree using the inoculation method above, with an inoculum volume of 3 μl. Each pseudo-replicate within a block was inoculated with the same source of inoculum. Conidial germination rate of the inoculum was determined as described above and ranged between 50-99% for different inoculation days.

The first measurements of lesion length were conducted 21 and 23 dpi for the top and bottom inoculation points respectively. Developing lesions were thereafter measured weekly for seven weeks

### Detached shoot experiment

The phenotyping of detached shoots was repeated in two years, with three replicate shoots/year. For each experiment, one-year dormant shoots were collected from trees of each of the members of the multiparental population. Shoots were collected from three different trees, to avoid pseudo-replication. All shoots were 60 (±10) cm long. The shoots were wrapped in damp tissue paper and stored at 4° C until the start of each experiment, when they were placed in wet floral foam (Oasis®) in large trays. The shoots were regularly supplied with water but not nutrients.

The experiments were carried out between February-May in year 1 (2017) and January-April in year 2 (2018) in a glasshouse compartment. Light and relative humidity were controlled in the same manner as for the potted tree experiment. Throughout the experiment the shoots were refreshed by cutting off approximately a centimeter at the bottom-ends with secateurs.

The inoculations of the two experiments were conducted as per Gómez-Cortecero et al. (2016) with a few modifications. Two leaf buds, the eighth and the fourteenth counting basipetally, were inoculated per shoot. An inoculum volume of 3 μl was pipetted onto each of the two leaf-scar wounds. Due to low lesion development following inoculation in year 2, the leaf-scars were re-inoculated 28 days after the initial inoculation.

Lesion lengths were recorded weekly, using digital calipers, after the first symptoms appeared. The lesion lengths were measured in six assessments in year 1 (between 23 and 58 dpi) and in seven assessments in year 2 (between 27 and 70 dpi).

### Phenotypic data

The area under the disease progression curve (AUDPC) was calculated for each genotype in the potted tree and detached shoot experiment using the R package ‘agricolae’ (de Mendiburu & Yaseen, 2020), whereas lesion size at five, eight and eleven mpi were used for the analysis of the field experiment. Cumulative data were used for the potted tree and shoot experiment rather than individual time-points as these experiments were carried out within a limited time-frame, and no differences were therefore expected between time-points.

Spatial corrections were assessed for the field and potted tree experiment using the package ‘SpATS’ in R (version 4.0.4; R Core Team, 2013) to remove variation due to tree position (Rodríguez-Álvarez et al., 2018). In the SpATS model, genotype and replicate were included as fixed effects and spatial coordinates within the field/glasshouse compartment (row and column) were included as random effects. BLUE for each genotype were thereafter generated using the SpATS model. BLUEs for the detached shoot experiment were obtained with the R package ‘lme4’ (Bates, Mächler, Bolker, & Walker, 2015) with genotype, year and lesion position as fixed effects, while tray nested within replicate were included as random effects. Broad sense heritability (H^2^) was calculated in SpATS. All data were either log or arcsin transformed before BLUEs were estimated. Back-transformed values were used for the analysis in FlexQTL for all phenotypes except %HTA and %CB, which remained arcsin transformed due to being skewed towards 100%.

Principal component analysis was performed using the function ‘prcomp’ in R, using scaled and centred BLUEs for each phenotypic trait.

### Genotypic data

DNA was extracted from flash-frozen leaf tissue of all genotypes in the multiparental population, including their parents and progenitors (Fig. 5). The DNA was extracted using EconoSpin® All-In-One Silica Membrane Mini Spin Columns (Epoch Life Science) according to the protocol for the DNeasy plant mini kit (Qiagen). The buffers used for extraction were according to Lamour & Finley (2006).

The population was genotyped on the Illumina Infinium® 20k SNP array (Bianco et al., 2014). The genotypes for each marker were assigned using GenomeStudio Genotyping Module 2.0 (Illumina). A subset of SNPs were selected after filtering by the software ASSisT (Di Guardo et al., 2013) and based on the absence of null-alleles in a previous set of 25 mapping populations and 400 pedigreed cultivars and breeding selections studied in the EU FruitBreedomics project (Laurens et al., 2018). SNP data curation and haplotype assignment were carried out according to Vanderzande et al. (2019). Approximately 6,000 SNP markers passed the quality control and were used to form 1083 haploblocks, distributed with 1 cM spacing across all chromosomes.

### QTL analysis

The QTL analysis was performed through Markov chain Monte Carlo (MCMC)-based Bayesian approach as embedded in the software FlexQTL™ (www.flexqtl.nl) as described by Bink et al. (Bink et al., 2002; 2008; 2012; 2014). The analyses were conducted using haplotype data, phenotypic BLUEs and a consensus genetic linkage map based on 21 full-sib families (Allard et al., 2016). Each QTL analysis had a MCMC chain of 200,000 iterations with a thinning of 200. The effective sample size in the parameter file was set to 100. To ensure reproducible results, multiple FlexQTL™ runs were conducted. Additional runs were made with different settings for 1) starting seed, 2) allowed maximum number of QTLs included in models (5,10), 3) prior for expected number of QTLs (3,5). QTL effects were set to being additive with a normal prior distribution and a (co)variance matrix with a random, diagonal structure. QTL positions were reproducible across the different settings. Furthermore, all the runs converged, and had effective chain size >100 for all parameters (Bink et al., 2014). Results are shown from the FlexQTL™ run with 10 maximum QTL and a prior of 5 QTL.

Two times the natural log of Bayes factors (2lnBF) was generated by FlexQTL™ and used to determine the level of significance of a QTL, where a 2lnBF of >2, 5, and 10 indicates a positive, strong, or decisive presence of a QTL, respectively (Bink et al. 2008; 2012; 2014). QTLs are assumed to be true if they had at least strong evidence for one QTL against no QTL on that linkage group (LG) within one experiment, or if they had at least positive evidence in two or more independent phenotyping experiments.

The QTL intervals were defined as regions covered by a continuous set of 2 cM bin intervals with 2lnBF>2. QTL intervals are reported for any QTL with at least positive evidence.

The proportion of phenotypic variation explained by each assumed true QTL was calculated using FlexQTL™ output and the formula: h^2^ = V QTL/V P, where V QTL = additive variance of a QTL and V P = total phenotypic variance.

### Haplotype analysis

The effect of single haplotypes on the canker phenotype at different time-points was estimated as follows. The size of haploblocks was reset in FlexQTL so that no recombinations occurred within assigned haploblocks in the multiparental population. The significance of haploblocks within the QTL regions was assessed in the software ASReml-R Version 4 (VSN International Ltd) to compare haplotype effects across time-points. A REML-analysis with the following model was used to test the significance of QTL-haploblocks:

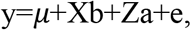

where y is a vector of observed BLUEs, X is the incidence matrix of fixed effects, b is a vector of fixed effects (in this case maternal/paternal haplotype), a is the vector of random effects (apple genotypes), Z is the design matrix of random effects and e is the residual error. The covariance structure for genotype effect was calculated using genomic information (i.e. N(0, Gσa 2)), where σ2a is the additive genetic variance and G the genomic relationship matrix. The genomic relationship matrix was constructed in R-package rrBLUP (Endelman, 2011). An inverse of the genomic relationship matrix was used in the asreml model.

The degree by which a haploblock can explain observed phenotypic variation was tested for each haploblock within the QTL region using a Wald statistic on the asreml-model. Maternally and paternally inherited haploblocks were tested separately. The haploblock with lowest *p*-value for each QTL was selected for further analysis.

The model above is expected to identify in which haploblocks the parental genotypes are segregating. However, it does not fully estimate the effect of haplotypes that may have been inherited from both maternal and paternal parents. Therefore, the number of copies of each haplotype (0, 1 or 2) within the selected haploblocks was included as a single fixed effect in the asreml-model described above. Only full-sib families were included in the haplotype analysis.

### Validation of haploblock effects

Disease phenotypes from 145 progeny resulting from the cross ‘Golden Delicious’ x ‘M9’ (GDxM9) were used to validate the effect of identified haploblocks. The cross progeny was phenotyped for resistance to *N. ditissima* in two separate detached shoot experiments and one potted tree experiment, as described above. Three replicate shoots were phenotyped in each of the two detached shoot experiments, whereas two replicate trees were used in the potted tree experiment. The lengths of canker lesions were measured in seven assessments in the detached shoot experiments (between 15 and 59 dpi) and in six assessments in the potted tree experiment (between 12 and 66 dpi). Marginal AUDPC means for each genotype was calculated using R-package ‘emmeans’ from a linear model with replicate and year as fixed effects. The genotyping and haplotype phasing of the GDxM9 family was performed as described for the multiparental population. A linear model was used to test the significance of each haplotype on AUDPC for each haploblock. A multi-QTL linear model with all of the significant haplotypes was thereafter fitted for both experiments to estimate variance attributed to each haplotype.

## Supporting information

Description of supplementary data

## Acknowledgments

The authors would like to gratefully thank Eric van de Weg (Wageningen University and Research) for advice on the analysis in FlexQTL as well as critically reviewing the manuscript. We would like to acknowledge the staff in the NIAB EMR farm and glasshouse department for assistance and support. A special thanks to Graham Caspall (Farm Manger, NIAB EMR) for all advice on the field experiment. This work was funded by the Biotechnology and Biological Sciences Research Council (BBSRC BB/P000851/1).

## Contributions

RJH and AK devised the study. AK performed the experimental work with input from AGC and CFN. AK performed all data analysis. All authors conceived and drafted the manuscript.

