## Supplementary material for "Identification of novel genetic regions associated with resistance to European canker in apple": Description of supplementary data

### Supplementary figures

**Supplementary figure 1A-1H.** Phenotypic distributions for each European canker phenotype by family.

**Supplementary figure 2**. Best Linear Unbiased Estimates (BLUEs) of parents and standard varieties**.**

**Supplementary figure 3**. Scree plot from the principal component analysis of European canker phenotypes

**Supplementary figure 4**. Phenotypic distributions for progeny of 'Golden Delicious' x 'M9'

### Supplementary tables

**Supplementary Table 1.** Mean and standard error for European canker phenotypes

**Supplementary Table 2.** Broad sense heritability for resistance to European apple canker in the multiparental population.

**Supplementary Table 3.** Summary of the selected haploblocks within QTL-regions

**Supplementary Table 4.** Haploblock alleles from parents segregating at QTL locus

**Supplementary Table 5.** Mean, standard error and significance of haploblock alleles in 'M9' x 'Golden Delicious' cross progeny
