## Supplementary figures for "Identification of novel genetic regions associated with resistance to European canker in apple"

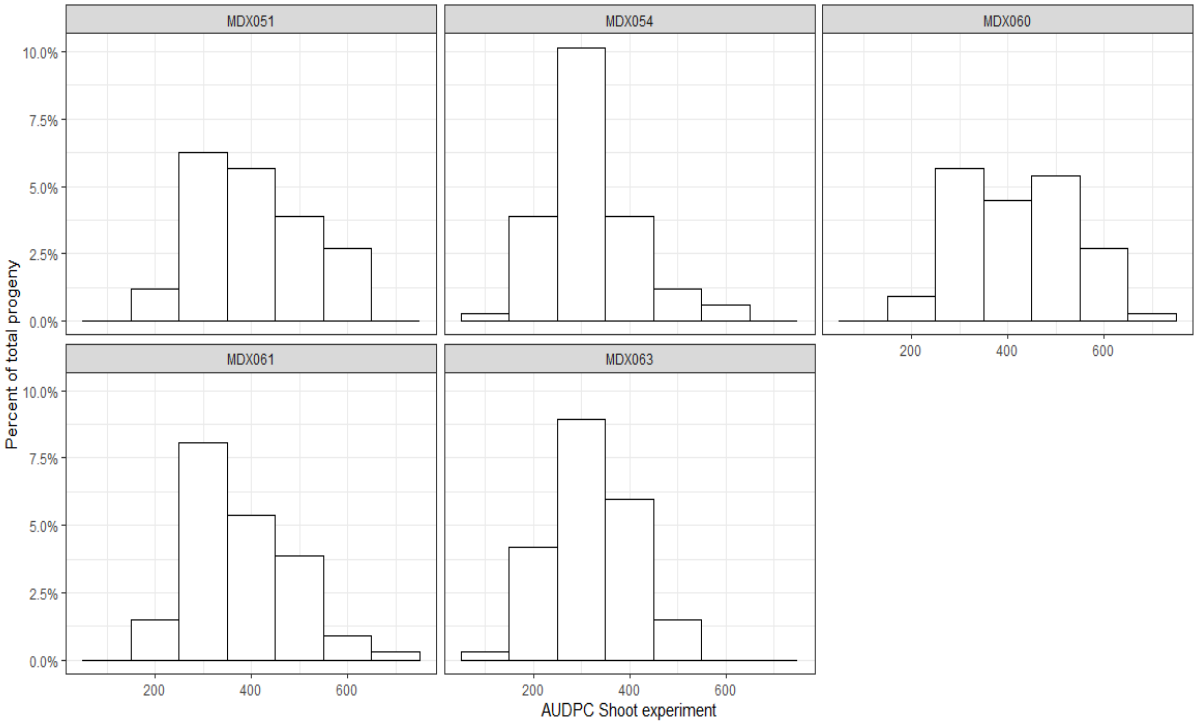


***Supplementary figure 1A***. Phenotypic distributions for susceptibility to European canker in the shoot experiment. Distribution shown by family.

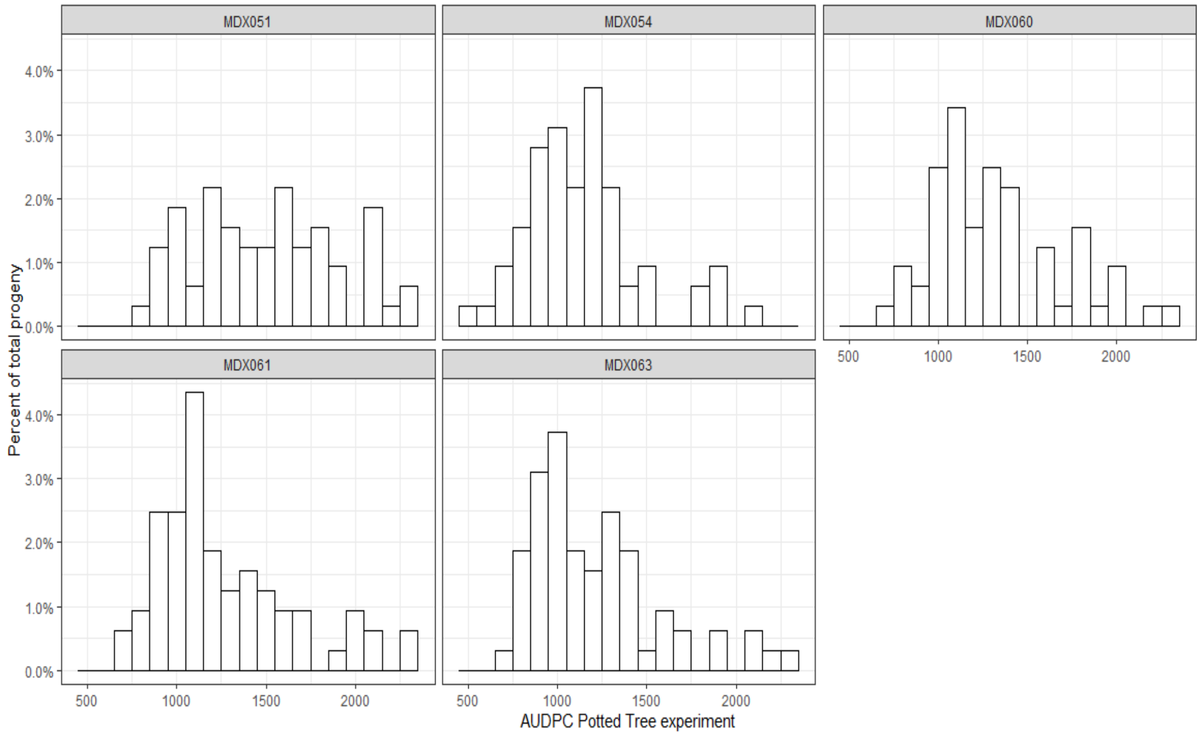


***Supplementary figure 1B.*** Phenotypic distributions for susceptibility to European canker in the potted tree experiment. Distribution shown by family.

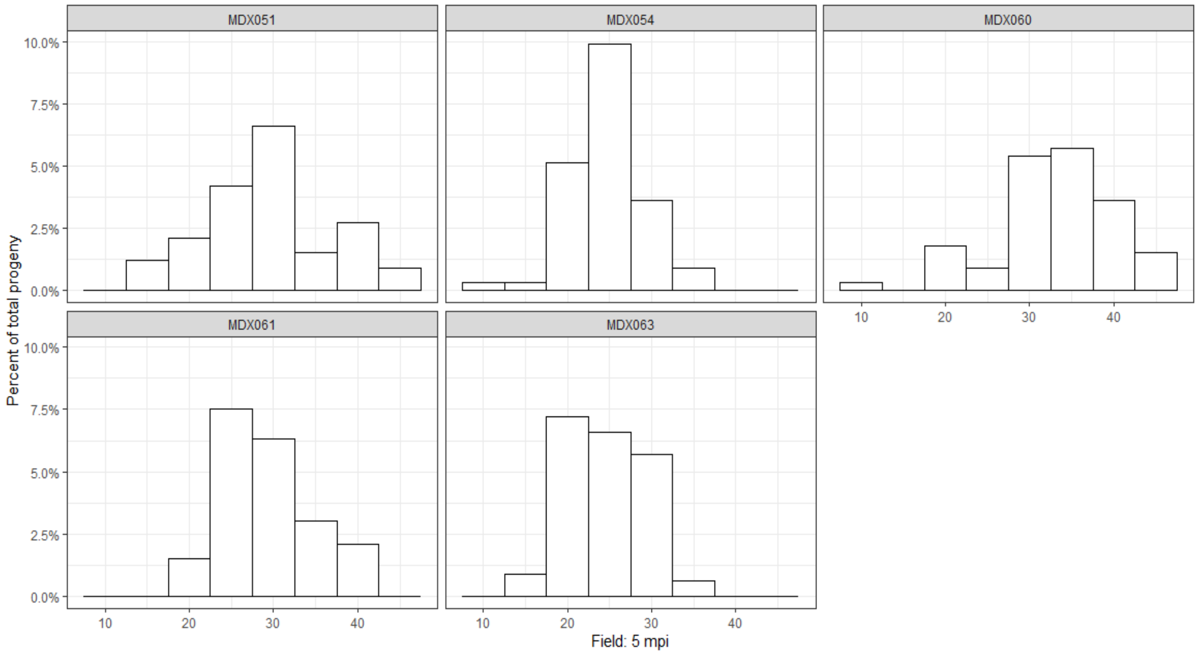


***Supplementary figure 1C.*** Phenotypic distributions for susceptibility to European canker in Field 5 mpi. Distribution shown by family.

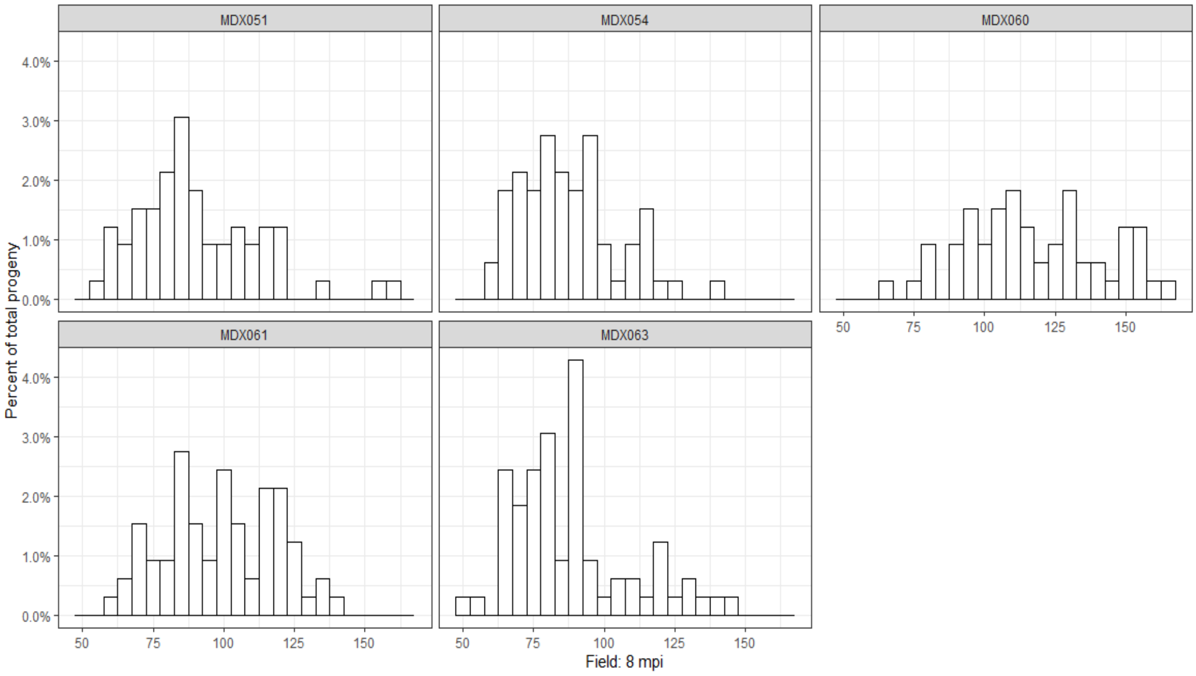


***Supplementary figure 1D.*** Phenotypic distributions for susceptibility to European canker in Field 8 mpi. Distribution shown by family.

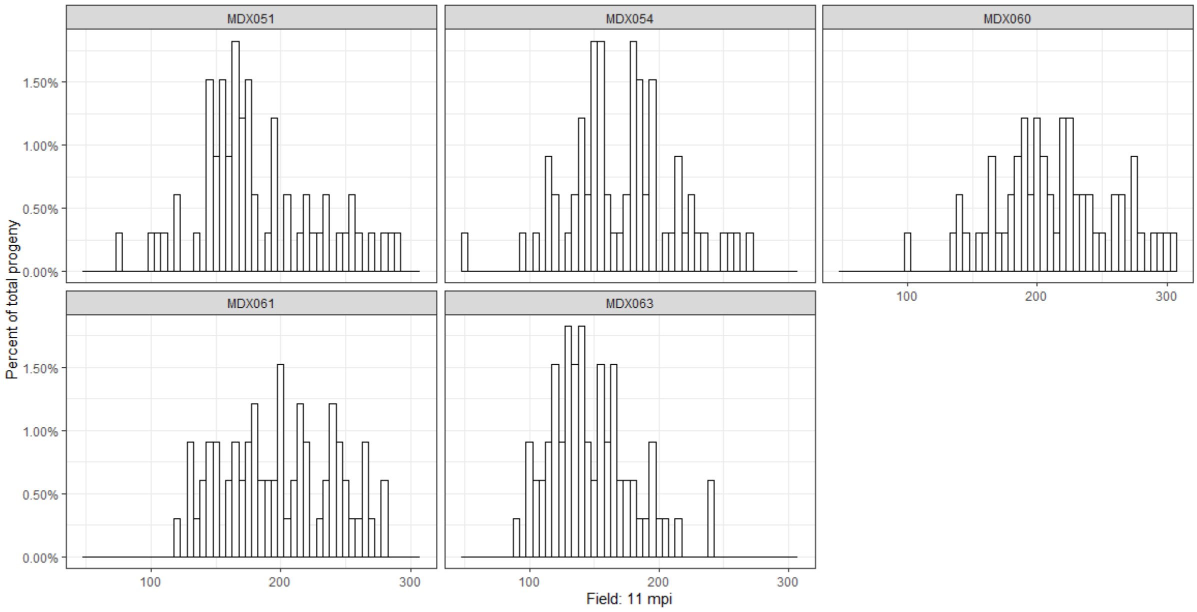


***Supplementary figure 1E***. Phenotypic distributions for susceptibility to European canker in Field 11 mpi. Distribution shown by family.

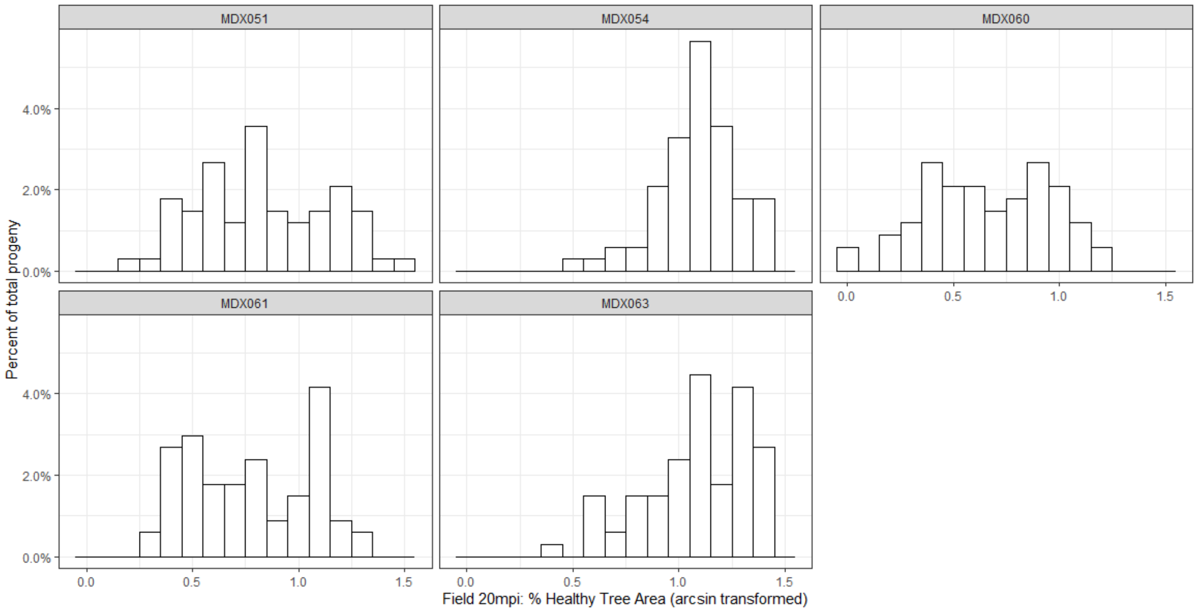


***Supplementary figure 1F***. Phenotypic distributions of arcsin transformed data for susceptibility to European canker in Field 20 mpi: % Healthy Tree Area. Distribution shown by family.

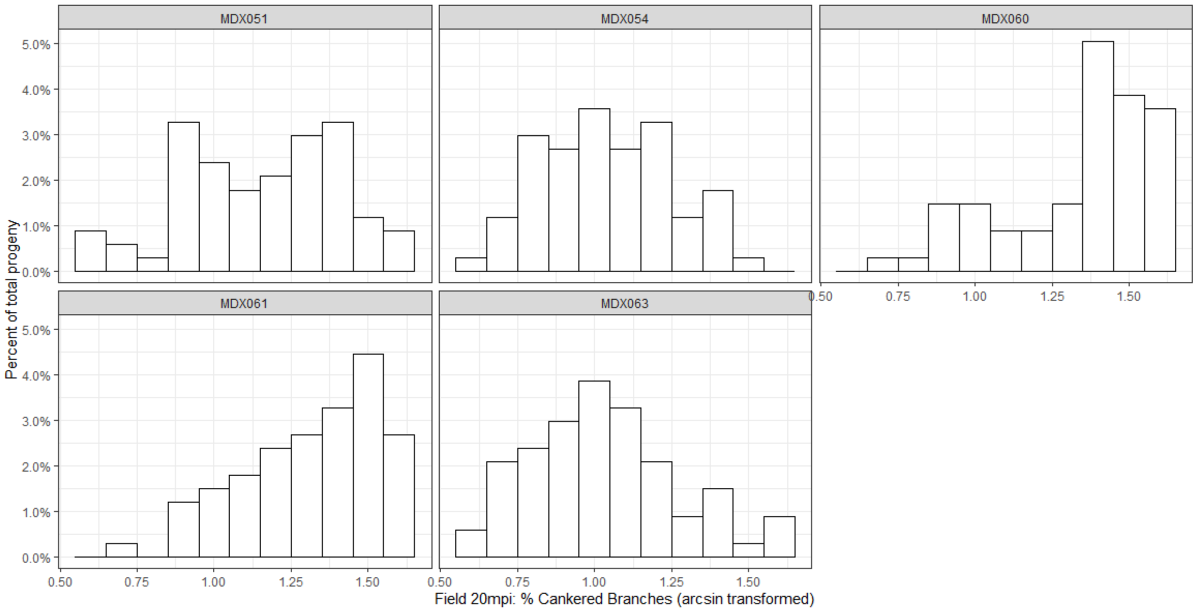


***Supplementary figure 1G***. Phenotypic distributions of arcsin transformed data for susceptibility to European canker in Field 20 mpi: % Cankered Branches. Distribution shown by family.

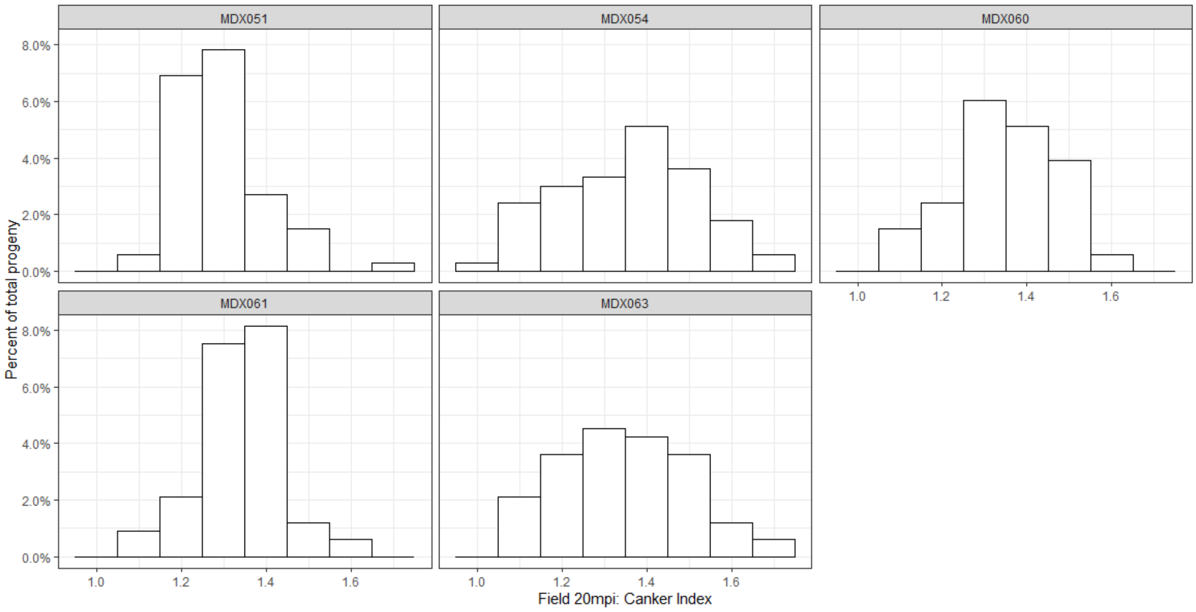


***Supplementary figure 1H.*** Phenotypic distributions data for susceptibility to European canker in Field 20 mpi: Canker Index. Distribution shown by family.

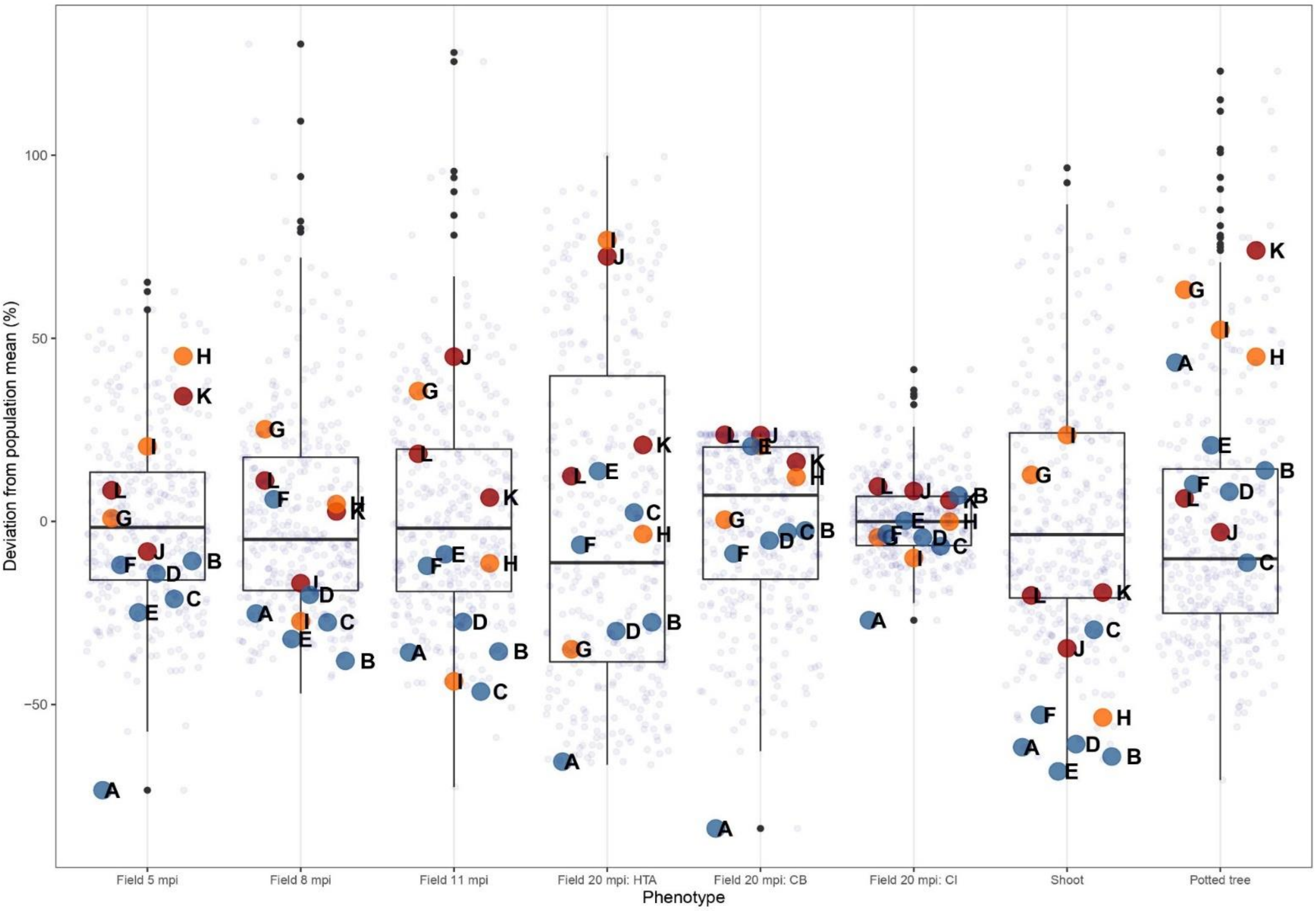


***Supplementary figure 2***. Best Linear Unbiased Estimates (BLUEs) of the multiparental population for each European canker phenotype. The data are shown as percent deviation from population mean, without the removal of outliers. Parents and standards are highlighted in colour, where the codes A-L refer to **(A)** seedling of *Malus robusta,* **(B)** Santana, **(C)** Jonathan, **(D)** Golden Delicious, **(E)** Elstar, **(F)** EM-Selection-1, **(G)** Aroma, **(H),** EM-Selection-2, **(I)** Cox Orange Pippin, **(J)** EM-Selection-4, **(K)** Gala, **(L)** EM-Selection-4. The inverse of %HTA is shown.

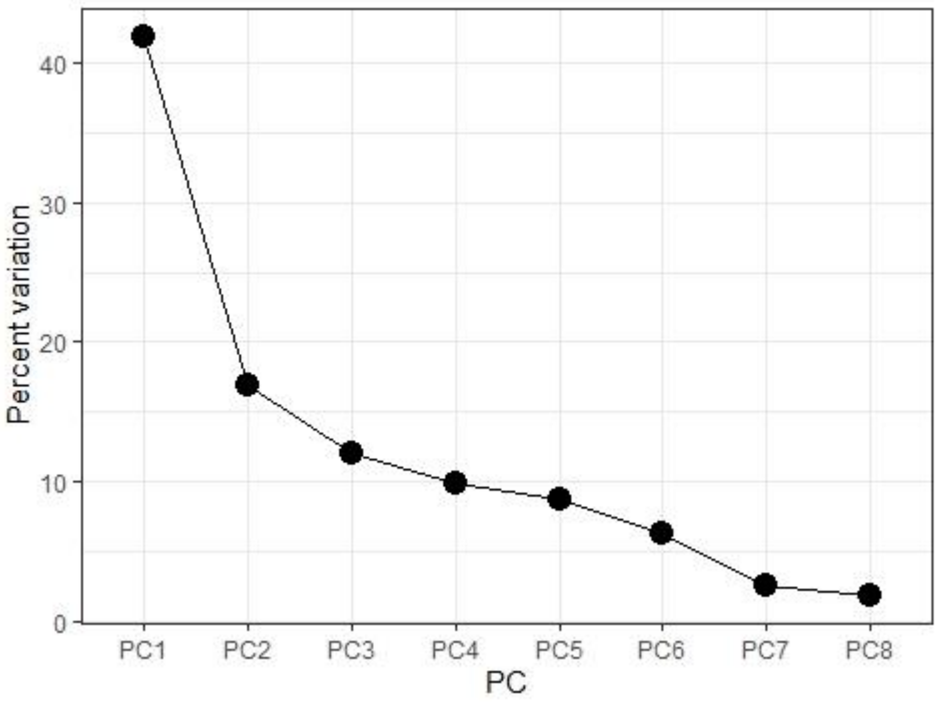


***Supplementary figure 3***. Scree plot from the principal component analysis (PCA) of the European canker phenotype data.

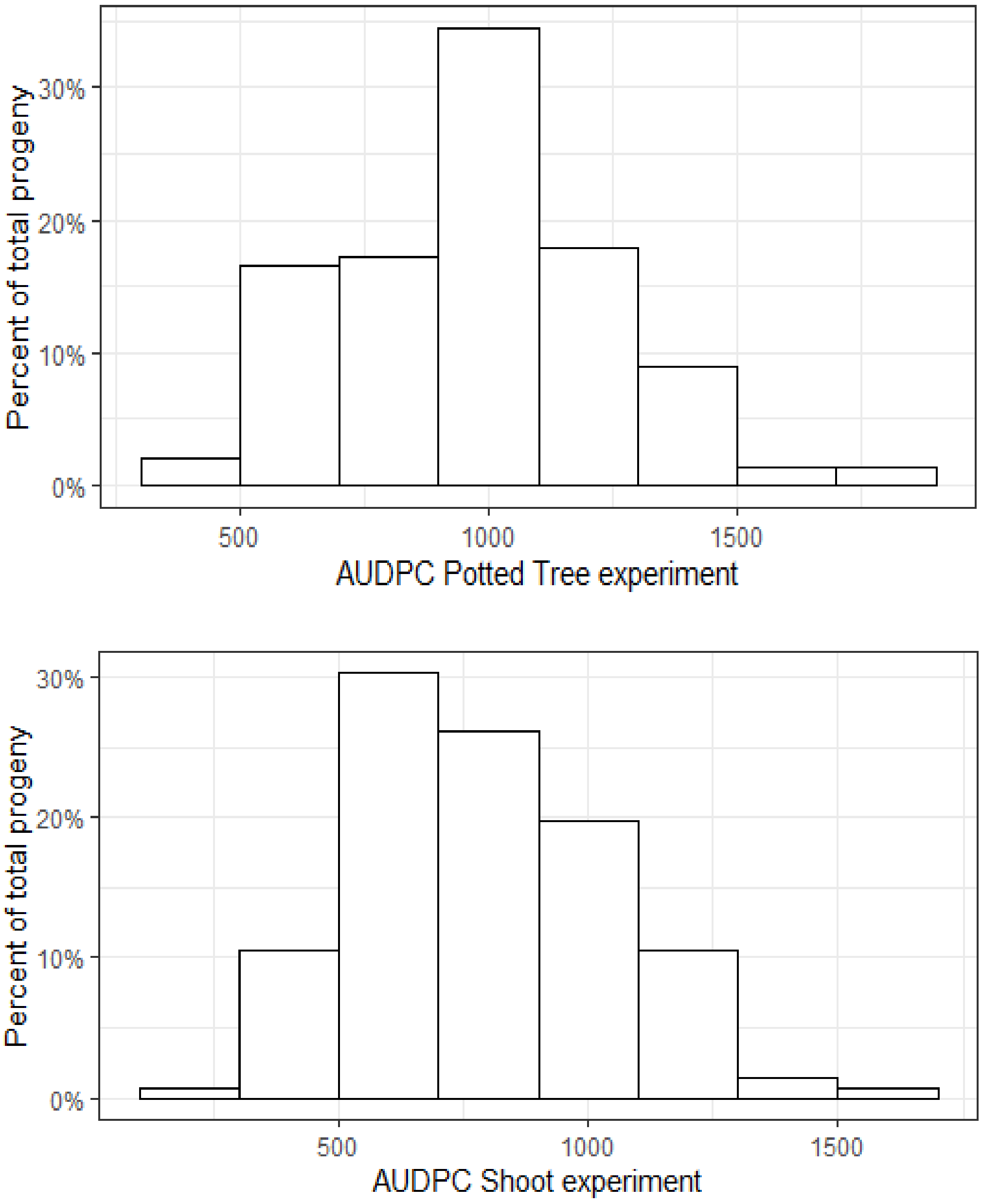


***Supplementary figure 4***. Phenotypic distributions for progeny of ‘Golden Delicious’ x ‘M9’ for susceptibility to European canker in a potted tree and a shoot experiment.
